# The linear correlation between genome size and the size of the non-transcribing region

**DOI:** 10.1101/2024.09.19.613789

**Authors:** Chen Zhang-Ren

## Abstract

**Background:** The genome sizes of organisms vary widely (C-value paradox). There are non-transcribing regions in the genome that neither encode proteins nor RNA entities. There are several hypotheses about the function of these regions: one suggests that they are unannotated functional areas, while another views them as genomic isolation zones that reduce mutations in coding regions.

**Method:** Statistical analysis was conducted on the transcribing regions (including areas annotated as genes and transcribed pseudogenes) and non-transcribing regions, protein-coding regions (Coding sequence, CDS), and genome sizes using annotation files from 63,866 species genomes in the NCBI RefSeq database.

**Results:** There is a significant linear relationship between the size of non-transcribing genomic regions and overall genome size across species, with varying proportional coefficients among different phyla (realms for viruses). As genome size increases, the proportion of non-transcribing regions gradually rises, eventually approaching a linear proportional limit, resembling one arm of hyperbolic functions. Eukaryotes show high linear correlation, with the highest in Streptophyta and the lowest in Apicomplexa. In eukaryotes, the size of the coding region increases with genome size, but the increasing trend diminishes (proportionally decreases). In non-eukaryotes, the size of the coding region maintains a linear relationship with genome size.

**Conclusion:** The size of non-transcribing region in species may be subject to some strict quantitative control mechanism, showing that genome and non-transcribing genome sizes increase proportionally with the expansion of the transcribing genome, indicating a strict balance between expansion and energy conservation. The proportion of non-transcribed genomes in eukaryotes is conservative (although the sequences are not), and the presence of non-transcribing genomes has significant implications for the evolution or survival of species. Thus, I propose a new hypothesis about the non-transcribing genome, that it is a space for generating new genes from scratch, and the different proportional coefficients among phyla are due to their different positions in energy transfer.

**Graphic Abstract:** 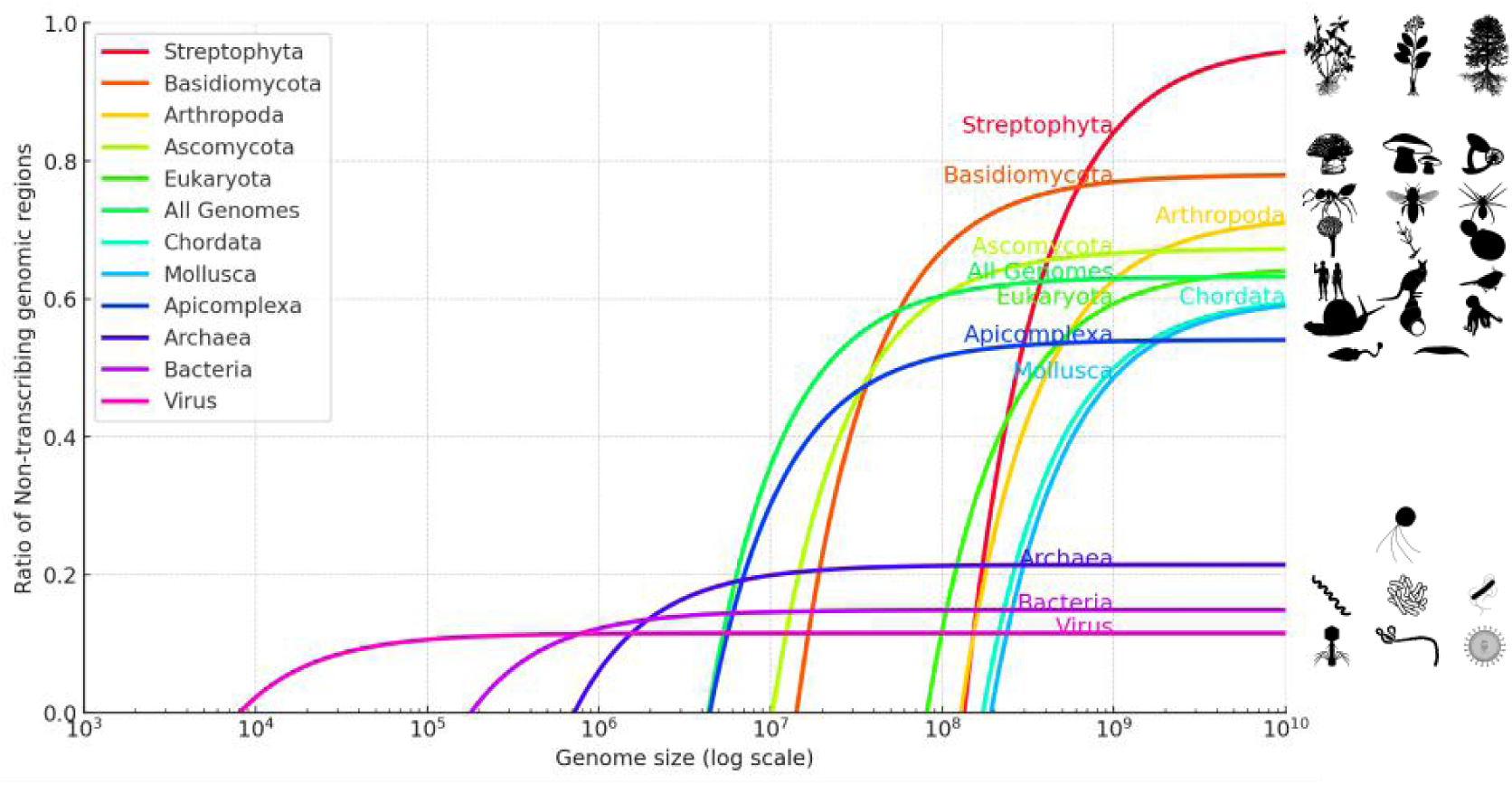

---

The genome size (C-value) and the size of the coding genome (G-value) vary significantly among different organisms^[1]^, a phenomenon known as the C-value paradox^[2]^ and the G-value paradox^[3]^. The genome consists of three types of DNA regions: (i) Coding regions: capable of encoding proteins or peptides, with proteins as the main functional entities; (ii) Non-coding but transcribing regions: unable to encode proteins or peptides but can be transcribed into RNA, with RNA as the main functional entities; (iii) Non-transcribing regions: neither translatable nor transcribable, with DNA as the main functional entity. In the human genome, 80-95% of the non-conservative parts are primarily composed of non-functional, decaying transposons^[4]^. Currently, researches on non-coding genome regions^[5]^, especially non-transcribed genome regions, is more insufficient than that on coding regions^[6]^. Regarding the function of the non-transcribed genome, hypotheses mainly include: (i) function unrevealed regions^[7]^, (ii) non-transcribing genome as non-functional areas, primarily serving to isolate genes and reduce mutations in functional areas^[8]^.

In studying the properties of the non-transcribed genome, I conducted a simple statistical analysis for genomes. First, I downloaded publicly available genome annotation files. Then, by calculating the size of the transcribing regions (excluding those annotated as genes and transcribed pseudogenes)/the non-transcribing regions and the size of the protein-coding regions (Coding sequence, CDS), I unexpectedly found a linear relationship between the sizes of non-transcribing or transcribing regions and the genome sizes across most species, with varying linear coefficients among different subgroup (phyla or realms). Therefore, I proposed a new hypothesis for the function of non-transcribing genomes: the non-transcribing genome regions serve as raw material for new gene formation from scratch (a “draft paper” hypothesis). The proportion of non-transcribing regions in a genome is an inherent characteristic of a species, governed by some yet fully elucidated precise regulatory mechanism, leading to genome expansion proportional to the transcription/non-transcription regions. Furthermore, based on the observation that the proportionality coefficient is highest in the Streptophyta phylum and lowest in the Apicomplexa phylum, I proposed the “energy constraint hypothesis.”

## 1. Method

### Database

Utilizing RefSeq genome and annotation database of NCBI^[9]^, genome annotating data (gff3) up to May 2023 was downloaded, including 77,118 publicly annotated genomes; after deduplication, 63,866 were obtained. These files contain annotated information for the genome of species. Among these species, there were 1,554 eukaryotes. And for non-eukaryotes, the genome with the highest assembly quality was selected for analysis.

### Data Processing

Scripts were written in Python to process the GFF3 files, which were used to extract and calculate the size of all coding regions (CDS), transcribing regions (including genes and transcribed pseudogenes), and non-transcribing regions in each species’ genome. The total genome size for each species was also obtained.

### Taxonomy

The study employed the NCBI taxonomy^[10]^ for species classification, which distinguishes between formal and informal names. Formal names are declared based on rules laid down in four relevant codes of nomenclature (although other codes do exist). These are the International Code of Nomenclature for algae, fungi, and plants (ICNafp), the International Code of Nomenclature of Prokaryotes (ICNP) and the International Code of Zoological Nomenclature (ICZN). The viruses are governed by the International Code of Virus Classification and Nomenclature (ICVCN, also referred to as the ICTV Code). Informal names follow internal rules that are dictated by practical considerations outside of the Codes. For example, names lacking species epithets are commonly applied to GenBank records.^[10]^

### Data Analysis Tools

GraphPad Prism 7 software was used for linear regression analysis to explore the relationship between the size of the transcribing region and the total genome size. Some charts (Fig. 1-5) were created using Prism 7. Fig. 6 was produced using the Python Matplotlib library called by GPT-4.0. The left curve part of the graphical abstract was completed using the Python Matplotlib library called by GPT 4.0.

**Fig. 1.**
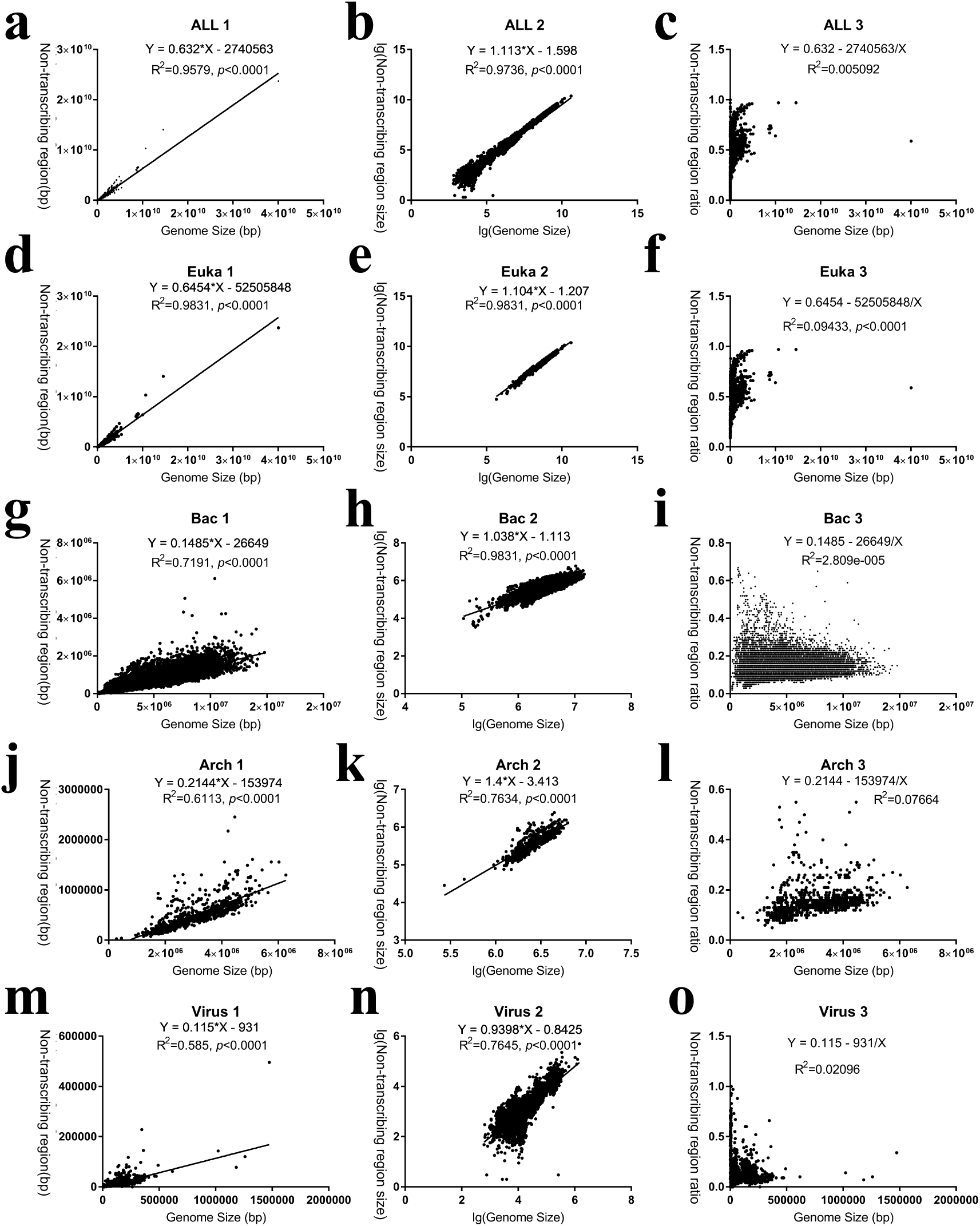
In eukaryotes, archaea, bacteria, and viruses, the size of non-transcribing regions is linearly correlated with genome size.

**Fig. 2.**
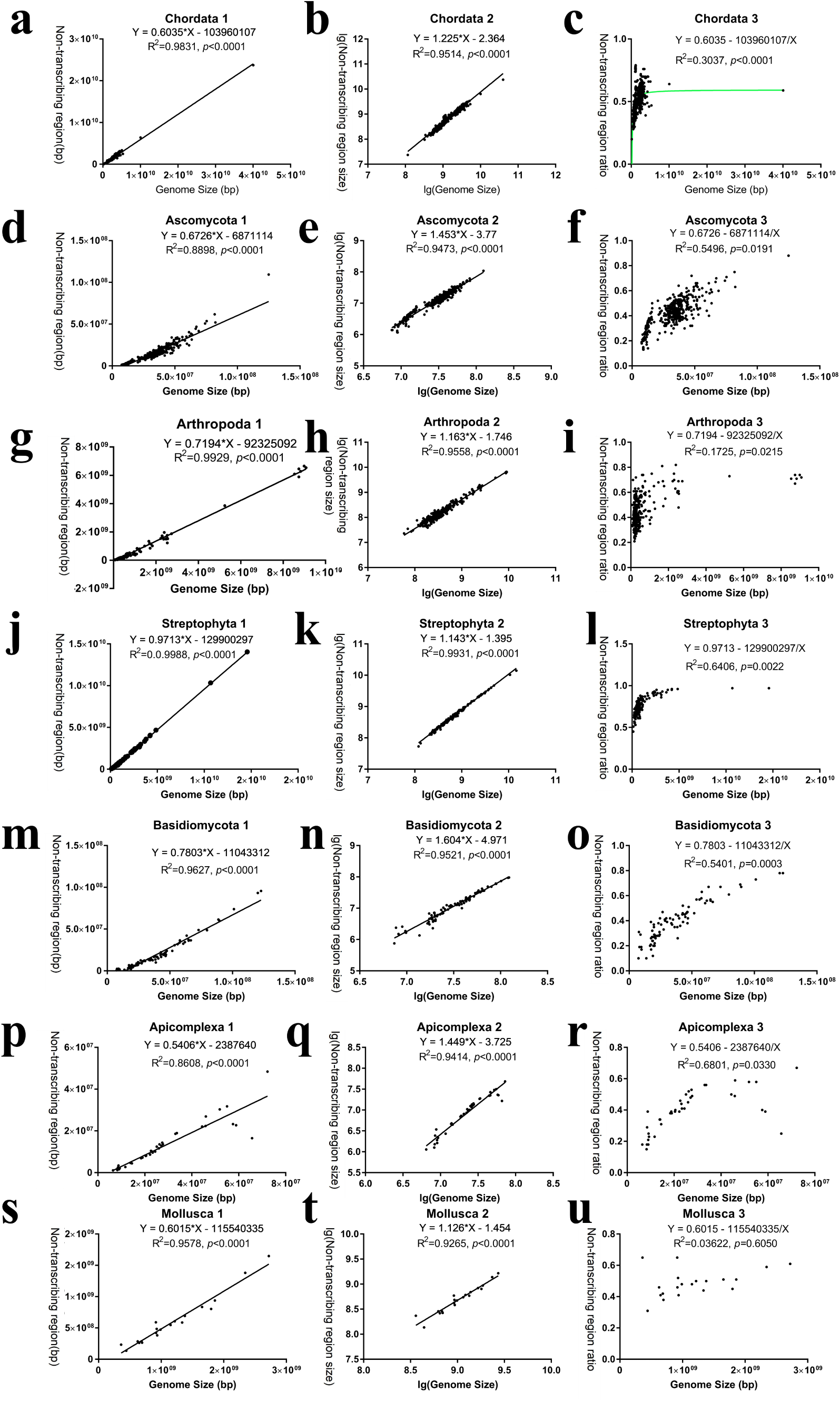
The linear relationships between non-transcribing region sizes and genome sizes vary across different phyla in eukaryotes.

**Fig. 3.**
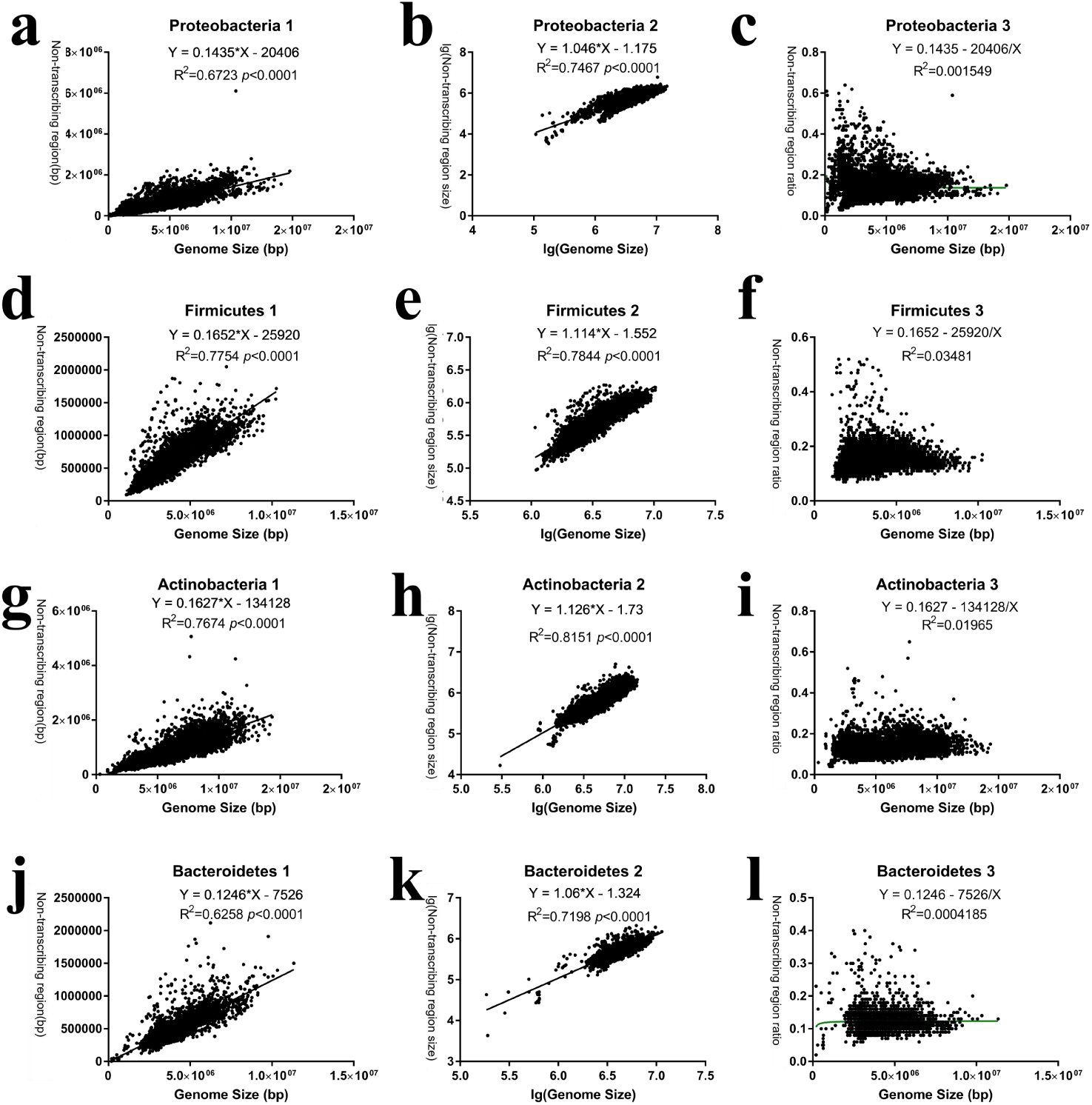
The linear relationships between non-transcribing region sizes and genome sizes in different phyla of bacteria.

**Fig. 4.**
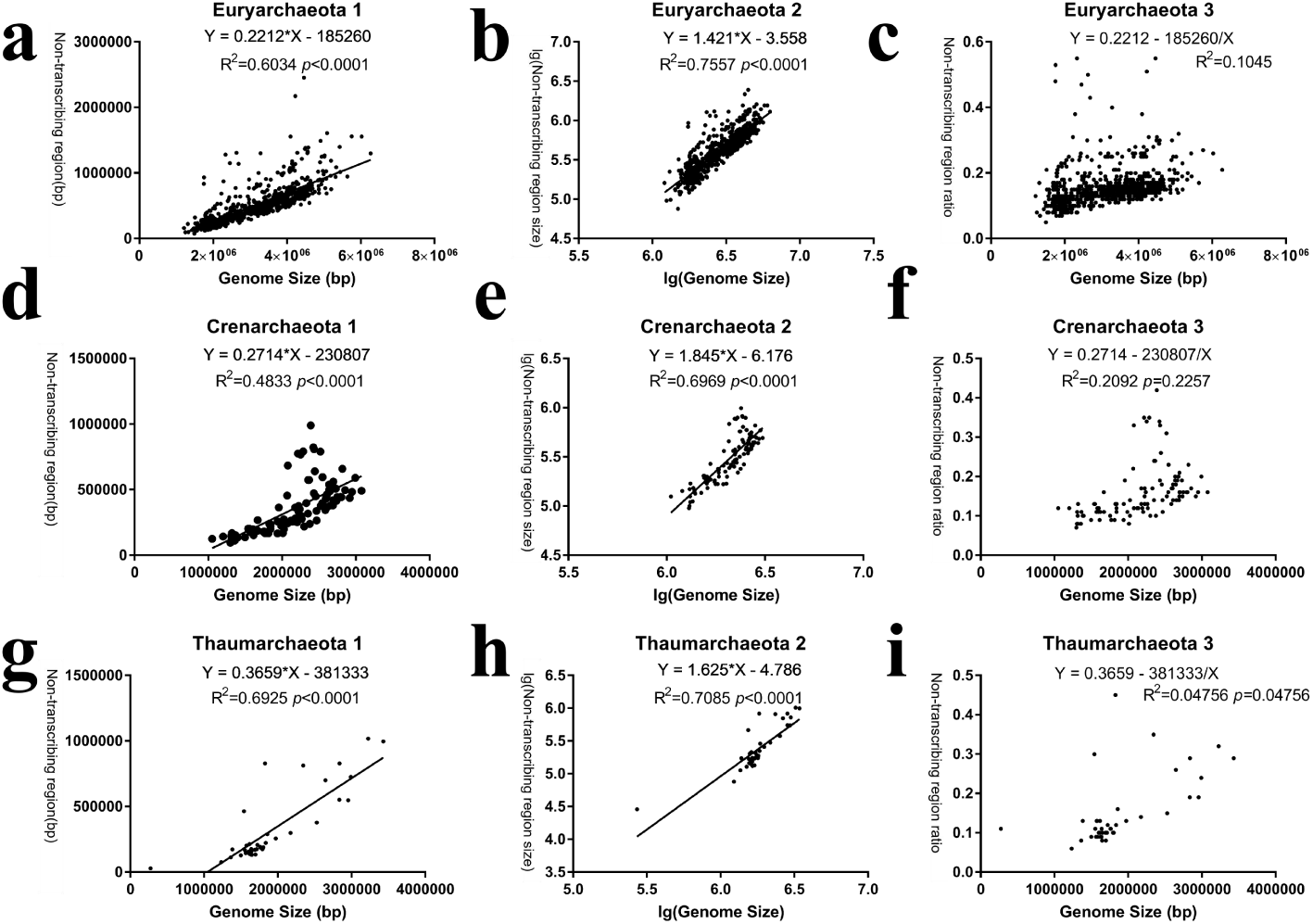
The linear relationships between non-transcribing region sizes and genome sizes in different phyla of archaea.

**Fig. 5.**
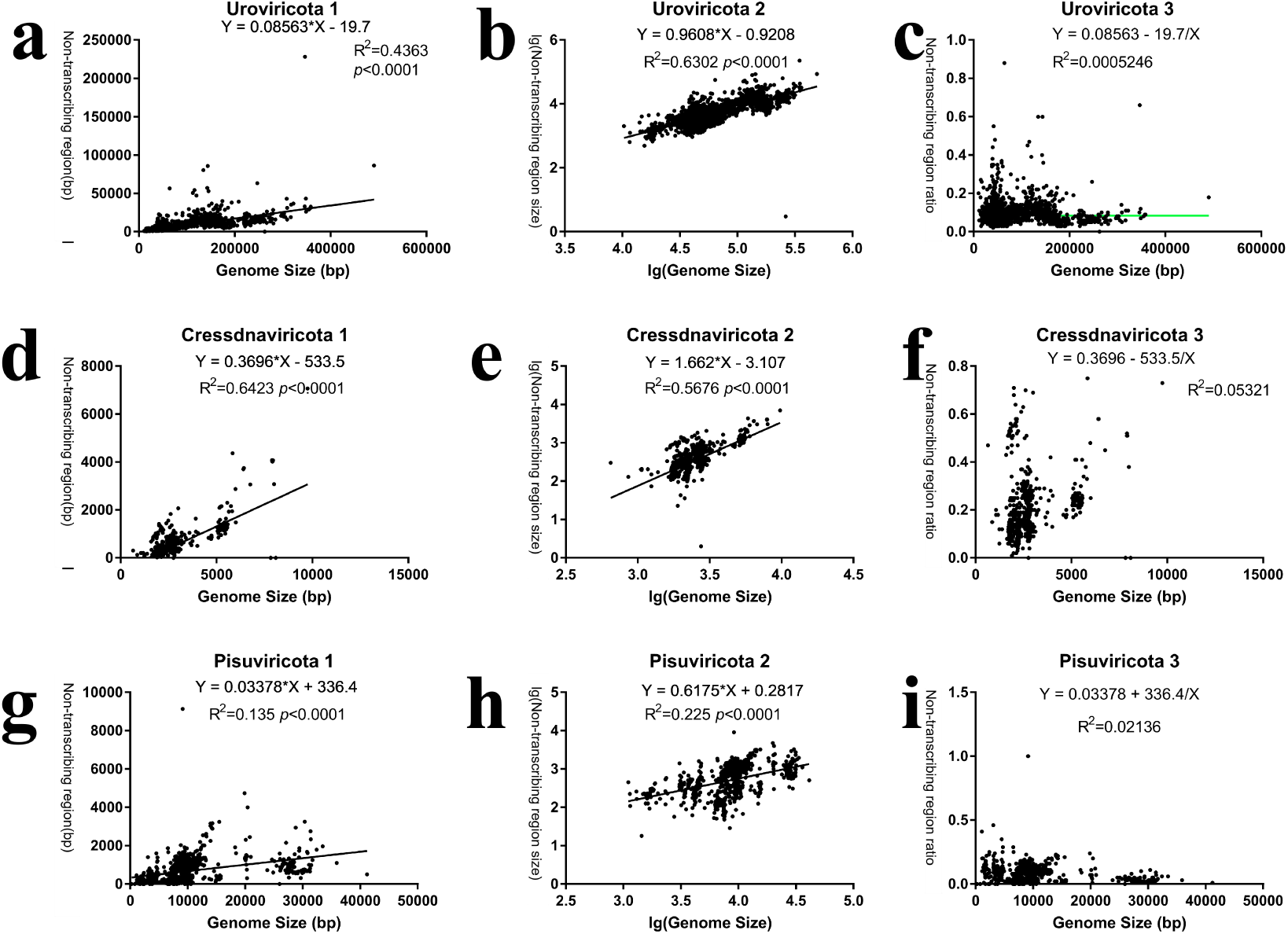
The linear relationships between non-transcribing region sizes and genome sizes in different realms of viruses.

**Fig. 6.**
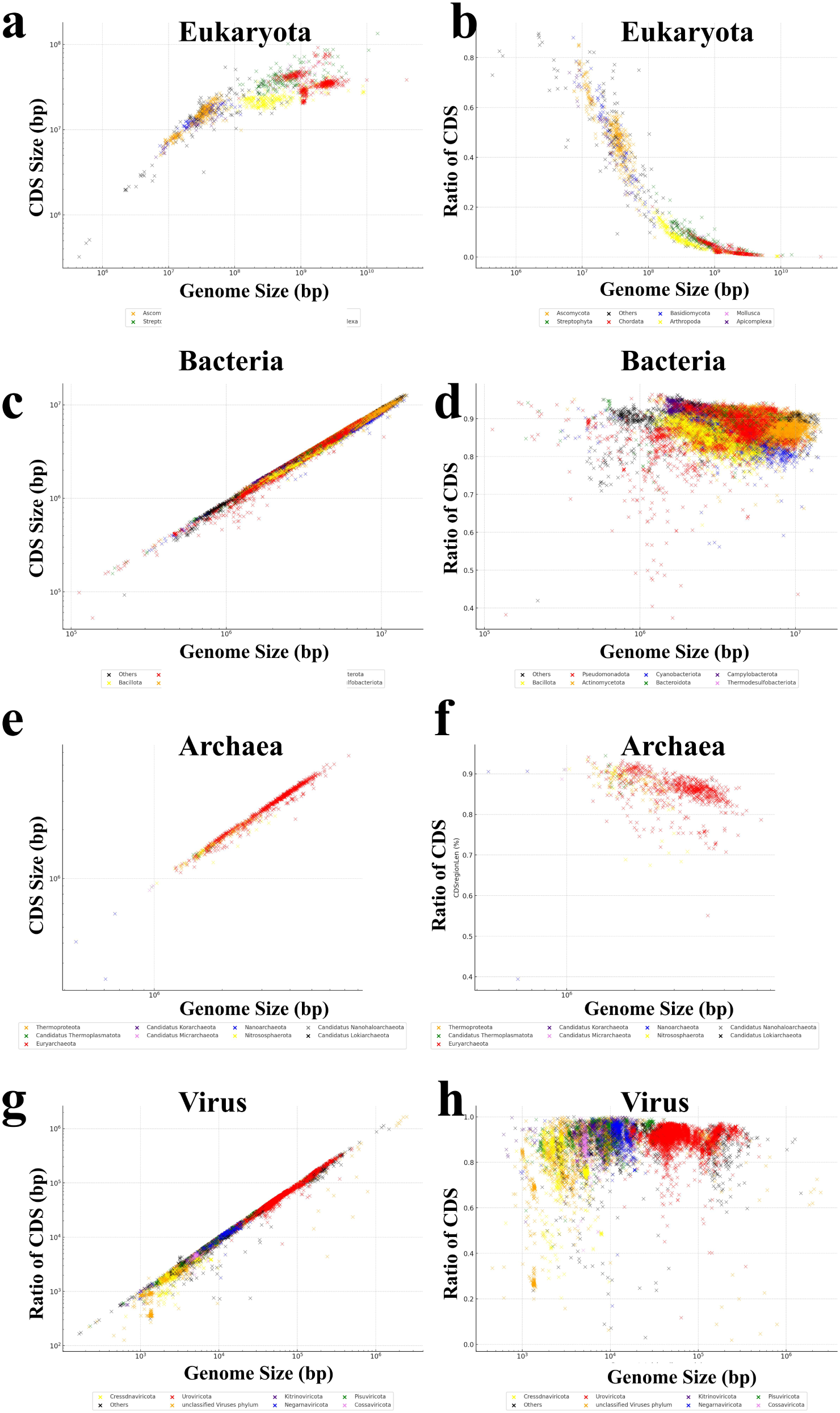
Trends and proportional changes of CDS region size of eukaryotes, bacteria, archaea, and viruses as genome size increases

## 2. Results

### 2.1. Proportional expansion of the genome: In eukaryotes, archaea, bacteria, and viruses, the size of non-transcribing regions is linearly correlated with genome size

A total of 77,118 publicly annotated genomes were downloaded from the NCBI RefSeq database. Of these, 13,252 genomes were removed due to incompleteness (e.g., genomes at scaffold or contig assembly levels) or redundancy. The remaining 63,866 genomes (including 1,554 eukaryotes, 52,294 bacteria, 917 archaea, and 9,101 viruses) were analyzed through their general feature format annotation files (.gff3 format files). I observed that the size of the non-transcribing genomic regions (N_Non-transcribing_) is linearly correlated with genome size (NGenome) (with linear p-values all less than 0.0001 Fig. 1), meaning that the size of the non-transcribing genomic regions (N_Non-transcribing_) increases proportionally with genome size (NGenome). These conform to Equation 1, where α and β represent the proportionality coefficient and bias term, both of which are constants greater than 0:

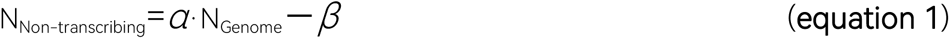

Since the proportion of the non-transcribing genomic region size (N_Non-transcribing_) to the genome size (N_Genome_) (Ratio of Non-transcribing Region, R_Non-transcribing_) is:

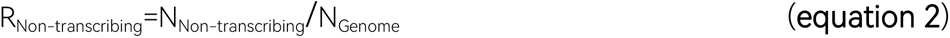

Thus, from Equation 1 and Equation 2, we can derive Equation 3:

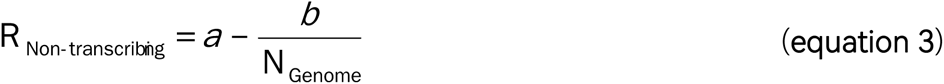

Equation 3 represents a branch of a hyperbolic function, consistent with the observations in Fig. 1cfilo and Fig. 2cfiloru: the proportion of the non-transcribing genomic region (R_Non-transcribing_) increases as the genome size (N_Genome_) grows, but the rate of increase gradually slows and eventually approaches a limiting value α.

Additionally, since genome size (N_Genome_) is composed of the size of the non-transcribing genomic region (N_Non-transcribing_) and the size of the transcribing genomic region (N_Transcribing_) (Equation 4):

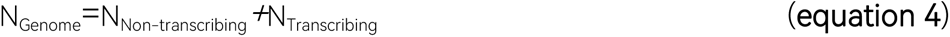

Therefore, the above linear relationship (Equation 1) can also be transformed into a linear relationship between the size of the transcribing genomic region (N_Transcribing_) and the genome size (N_Genome_) (Equation 5), as well as between the size of the transcribing(N_Transcribing_) and non-transcribing genomic regions(N_Non-transcribing_) (Equation 6):

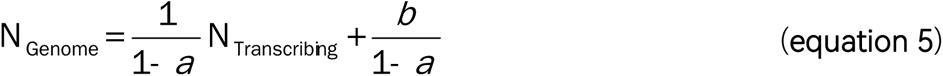

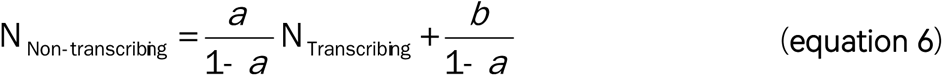

That is to say, if we consider the size of the transcribing genomic region (N_Transcribing_) as the initial changing value (i.e., when new genes are born), the genome size (N_Genome_) and the size of the non-transcribing genomic region (N_Non-transcribing_) will increase proportionally. For instance, with each 1 bp increase in the transcribing genomic region, the non-transcribing genomic region increases by an average of 2.56 bp in Arthropoda (resulting in a total genome increase of 3.56 bp), and by an average of 33.84 bp in Streptophyta (resulting in a total genome increase of 34.84 bp). I refer to this as the phenomenon of proportional expansion of the genome.

### 2.2. Proportional Coefficient Differences and Conserves: Variations in the Linear Relationship Between Non-transcribing Region Size and Genome Size Across Different Phyla/Realms, with High Conservation in Eukaryotes

In studying the linear relationship between the size of non-transcribing regions and genome size across eukaryotes, archaea, bacteria, and viruses, I observed that as genome size increases, the distribution range of non-transcribing region sizes diverges (Fig. 1). Therefore, I further investigated the linear relationships within different subgroups (phyla/realms, with “realm” specific to viral classification) of eukaryotes, archaea, bacteria, and viruses. After accounting for these subgroups, the divergence significantly decreased, and the fit between the non-transcribing region size distribution and a hyperbolic function improved.

As shown in Fig. 2-5 and Tab. 1, the linear relationship between non-transcribing region size and genome size varies across different phyla/realms. This variation is particularly noticeable in eukaryotes. For instance, the Streptophyta phylum has the highest proportionality coefficient (α value), followed by Basidiomycota, while Apicomplexa has the lowest. Additionally, the linearity is highest for Streptophyta (R²=0.9988). These findings suggest that the proportionality coefficient α between non-transcribing region size and genome size might be a conserved feature across different phyla/realms, with particularly high conservation in eukaryotes. This indicates that it is not the sequence of the non-transcribing regions (which contain many repetitive elements) that is conserved, but rather their quantity.

**Table 1.**
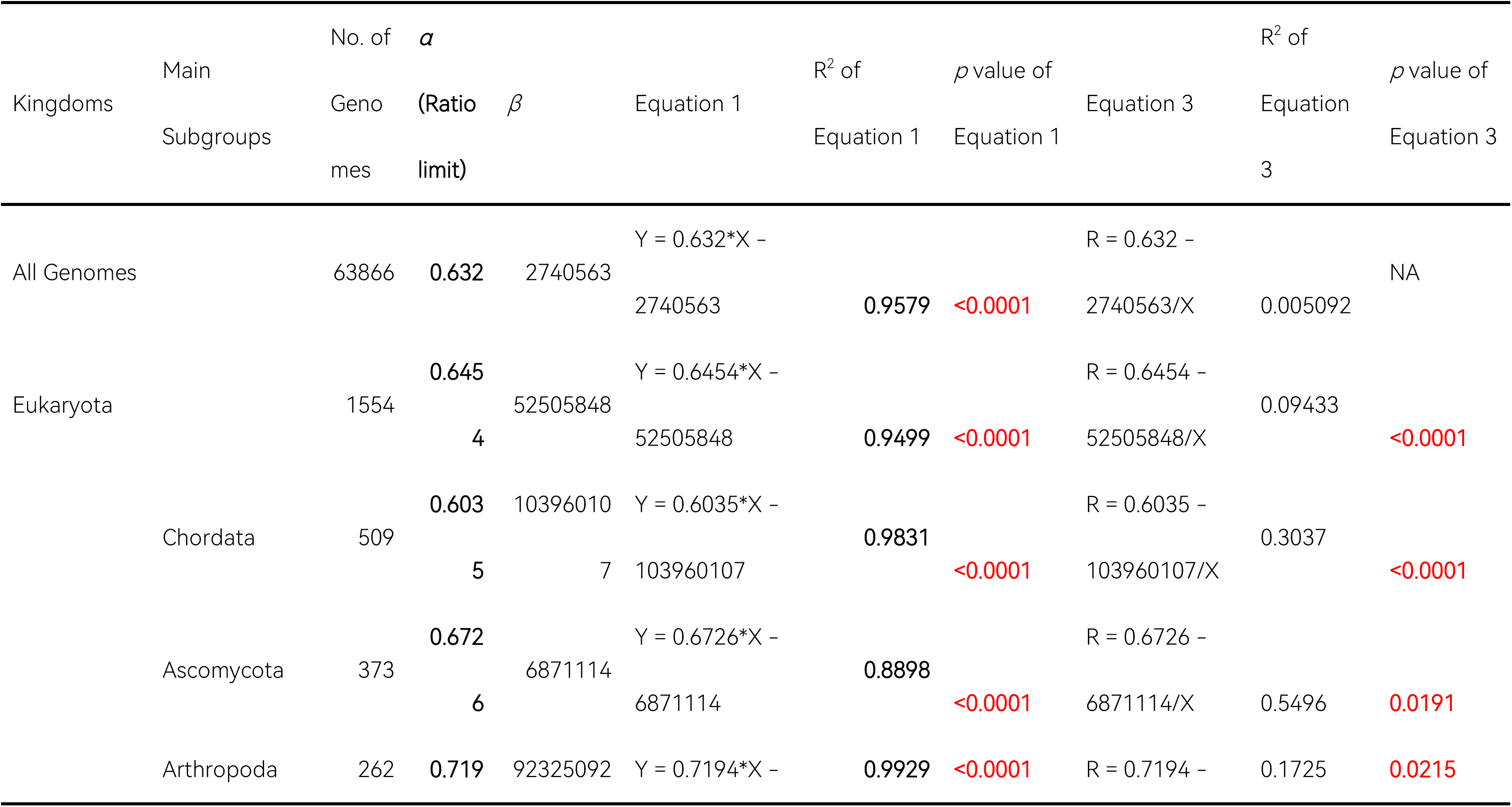
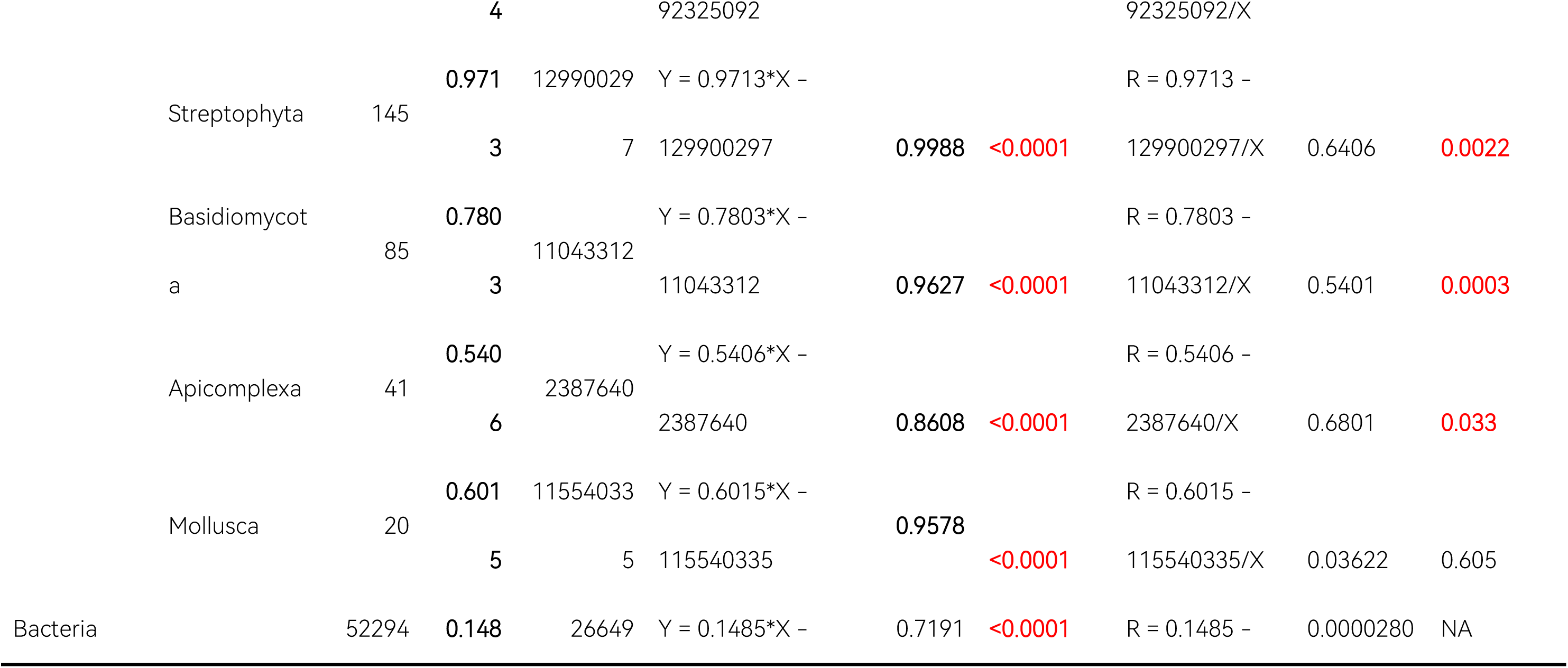
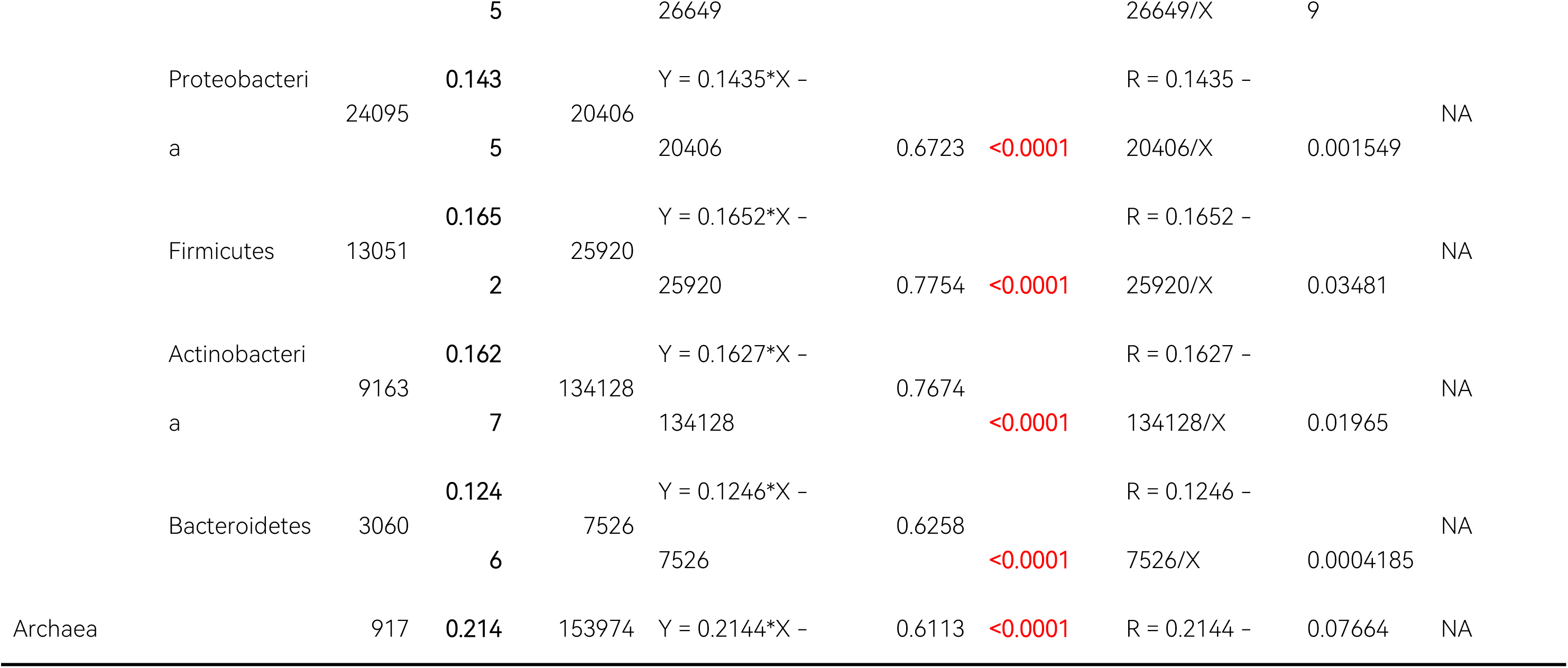
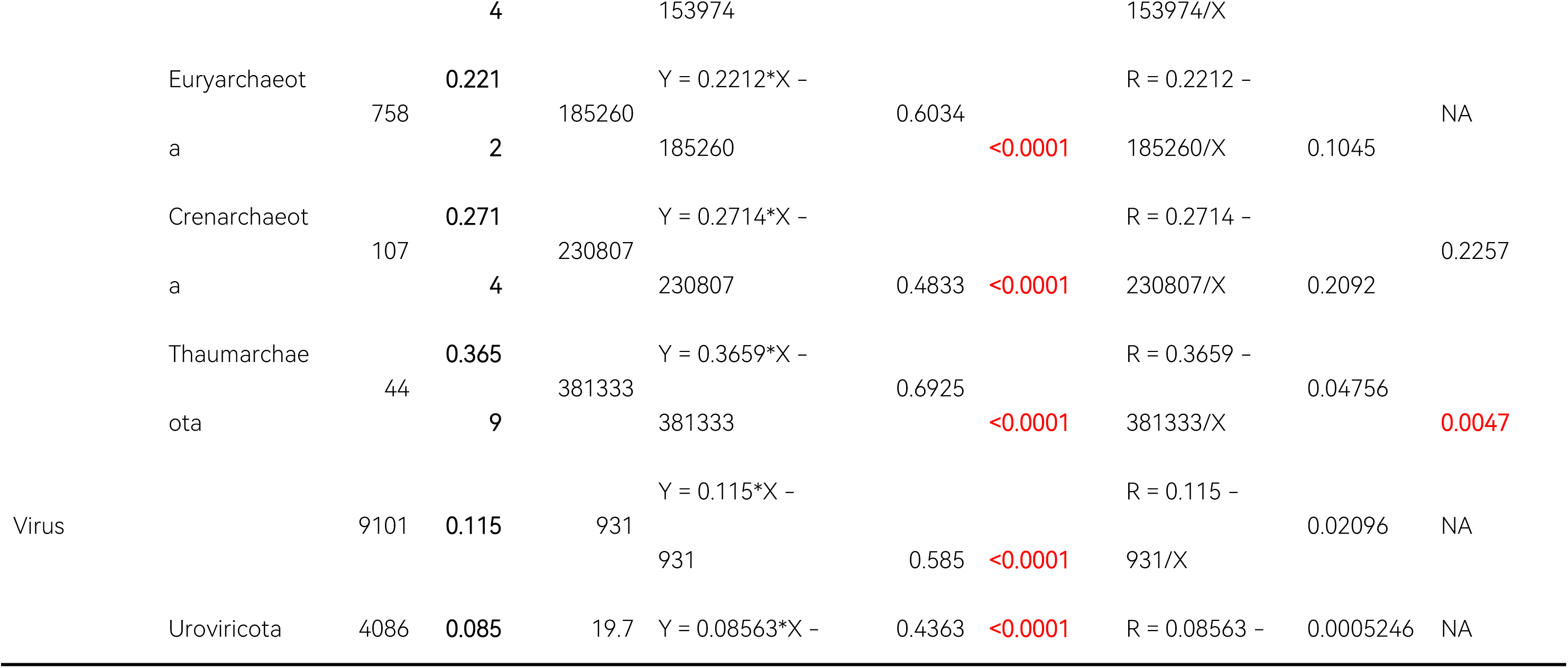
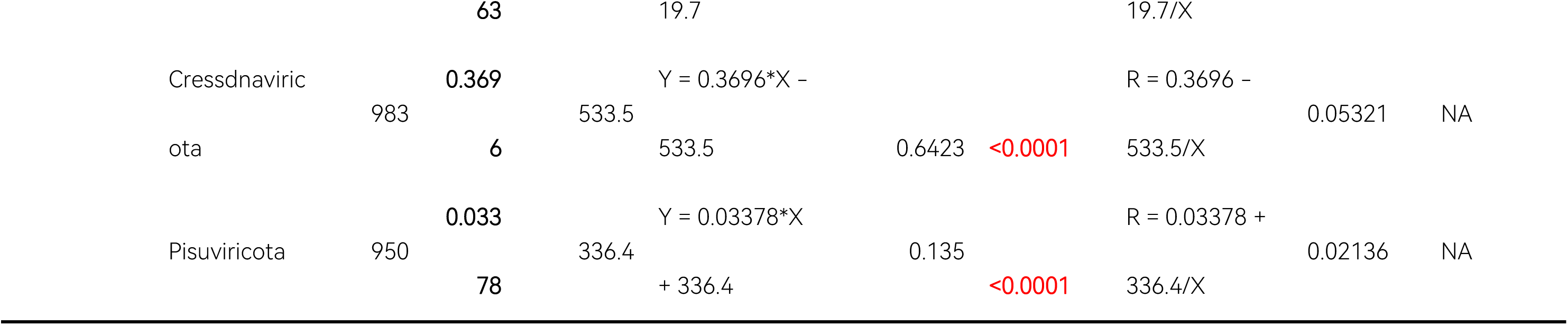
The linear relationship between the size of non-coding regions and genome size in different organisms.

For eukaryotes, the proportionality coefficients are as follows: Streptophyta = 0.9713, Basidiomycota = 0.7803, Arthropoda = 0.7194, Ascomycota = 0.6726, Chordata = 0.6035, Mollusca = 0.6015, and Apicomplexa = 0.5406. In contrast, for non-eukaryotes, the coefficients are much lower: Archaea = 0.2144, Bacteria = 0.1485, and Viruses = 0.115.

Why do these proportionality coefficients vary so widely across subgroups? I believe this may be related to the order of energy flow among organisms, and I propose the “energy constraint hypothesis” to explain this phenomenon (see Discussion).

### 2.3 In eukaryotes, the sizes of the coding sequence (CDS) region tends to increase with genome size, but this trend gradually slows down. In contrast, the size of the CDS region in non-eukaryotes generally maintains a linear relationship with genome size

To investigate the relationship between the size of the coding genome, which primarily functions by translating proteins, and the genome size of species, I extracted and analyzed the total length of the CDS (coding sequence) region for each species (excluding overlap). I also analyzed the ratio of CDS regions in genomes.

The results show a non-linear positive correlation between coding genome size and genome size in eukaryotes (Fig. 6a). When genome size is relatively small, there is a significant positive correlation between the two variables. However, as genome size continues to increase, the rate of increase in coding genome size gradually slows, indicating a saturation effect or diminishing marginal returns. Additionally, the proportion of the genome occupied by the coding genome rapidly decreases as genome size increases (Fig. 6b). This suggests that as the coding genome size increases in eukaryotes—corresponding to the emergence of new protein-coding sequences—the genome size increases non-linearly, with the slope gradually increasing. These results are consistent with the findings of Sebastian et al.^[11]^.

In contrast, for non-eukaryotes, bacteria, archaea, and viruses, there is a linear positive correlation between coding genome size and genome size, and the proportion remains relatively stable or decreases slightly (Fig. 6c-h). This suggests that as the coding genome size increases in non-eukaryotes—the genome size increases in roughly proportional linear increments.

## 3. Discussion

My quantitative analysis of genomes reveals a linear correlation between non-coding region size and genome size across eukaryotes, archaea, bacteria, and viruses. This indicates a phenomenon of proportional expansion in genomes. The linear relationship between non-coding region size and genome size differs among organisms from different phyla/realms, with eukaryotes exhibiting higher linear conservation. Furthermore, quantitative analysis on protein-coding regions shows that in eukaryotes, the size of the CDS region increases with genome size, but the rate of increase gradually slows. However, in non-eukaryotes, the size of the CDS region remains linearly correlated with genome size.

There are three parts of genome: (1) the coding genome, where proteins serve as the functional entities, (2) the non-coding, transcribing genome, which transfers into RNA, and (3) the non-transcribing genome, where DNA is the functional entity (Fig. 7). These results suggest that those regions are quantitatively related. This brings new perspectives on the functional interpretation of the non-transcribing genome: the non-coding genome is a functional component of the genome, where its sequences are non-conserved and unordered, yet the quantity is kept at a fixed proportion critical for the survival of the species. It does not expand chaotically due to transposon replication^[12]^ nor does it diminish due to DNA loss processes like programmed DNA elimination (PDE)^[13,14]^. These quantitative relationships may partially resolve the “C-value paradox,” but they also shift the focus to the “G-value paradox”^[15]^. Is there a pattern to the size of coding genomes in different species?

**Fig. 7.**
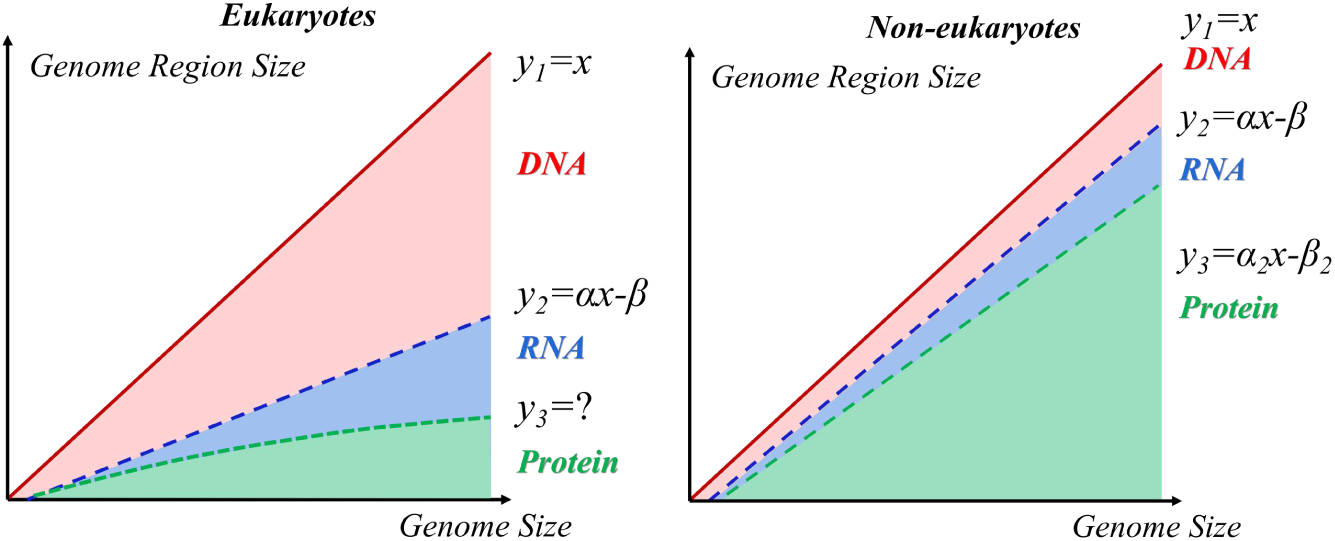
Diagrams for quantitative relationships between genome size, coding genome size, non-coding transcribing genome size, and non-transcribing genome size in eukaryotes and non-eukaryotes.

Previous hypotheses about the non-coding genome include the ENCODE hypothesis, which suggests that all non-coding sequences in the genome are functional but are simply uncharacterized functional components^[7]^. “Far from being ‘junk’, the DNA between protein-coding genes consists of myriad elements that determine gene expression, whether by switching transcription on or off, or by regulating the degree of transcription and consequently the concentrations and function of all proteins.”^[16,17]^ However, this hypothesis has been questioned, particularly when estimates of the functional genome are based on mutation rates. In 2017, Dan Graur developed a mathematical model based on human genome size, deleterious mutation rates, and fertility rates, concluding that perhaps only 8-14% of the human genome is functional, and this percentage likely does not exceed 15%.^[8]^ In 2014, Chris M. Rands estimated that no more than 8.2% of the genome is functional.^[18]^ This supports another hypothesis that the non-coding genome acts as a buffer, protecting the functional genome from mutations. “Cell biology may require a certain C-value, but most of the stretches of noncoding DNA that go to satisfying that requirement are junk (or worse, selfish).”^[19]^ However, neither of these hypotheses fully explains the strict linear results observed in this study.

Based on the above findings, I propose a new hypothesis: the non-transcribing genome serves as “draft material” for de novo gene formation^[20]^, providing the necessary space for new gene creation. The stochastic transcription of the non-transcribing genome (which I term “transcribing fluctuation”) leads to the emergence of unstable genes (which I refer to as “virtual genes”), which through natural selection may give rise to genes that improve fitness, becoming stably integrated into the genome as “real genes” and potentially leading to new protein-coding genes (Fig. 8). Eukaryotes, due to the difficulty of acquiring new genes by the mechanisms of horizontal gene transfer which is common among non-eukaryotes, rely more heavily on de novo gene formation from the non-transcribing genome. This explains why eukaryotes have larger and more conserved proportions of non-transcribing genomes compared to non-eukaryotes. This process, which creates new genes and proteins from nothing, is somewhat analogous to the generation of particles from vacuum fluctuations in physics, where the non-transcribing genome functions as the necessary “raw material” or “draft paper.” While retaining a large non-coding genome increases the likelihood of de novo gene formation (leading to new traits and increased fitness), it also consumes resources and energy and may pose disadvantages to the cell^[21,22]^. Thus, there is a balance between the expansion and reduction of the non-transcribing genome, which I believe is primarily determined by the organism’s position in the energy transfer chain within an ecosystem. As primary producers, organisms in the phylum Streptophyta (mainly plants and photosynthetic algae) occupy the initial position in the energy transfer chain and can retain the highest proportion of non-coding genomes. In contrast, organisms in the phylum Apicomplexa, which occupy a parasitic niche as consumers, have the least access to energy and therefore retain the lowest proportion of non-coding genomes among eukaryotes. I refer to this as the “energy constraint hypothesis” (Fig. 9).

**Fig. 8.**
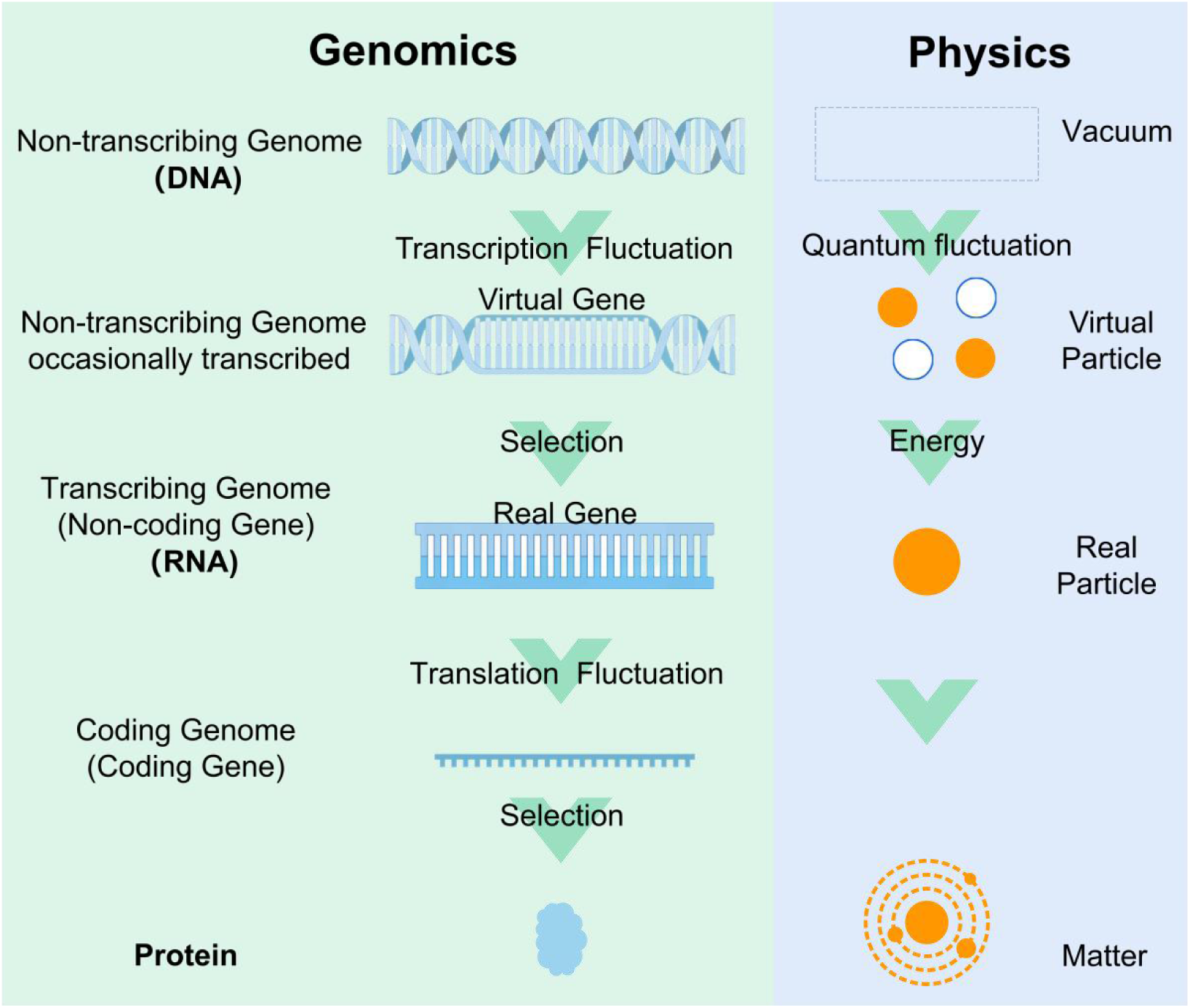
The “Draft Paper Hypothesis” of non-transcribing genome function (analogous to the creation of new particles from vacuum fluctuations in physics).

**Fig. 9.**
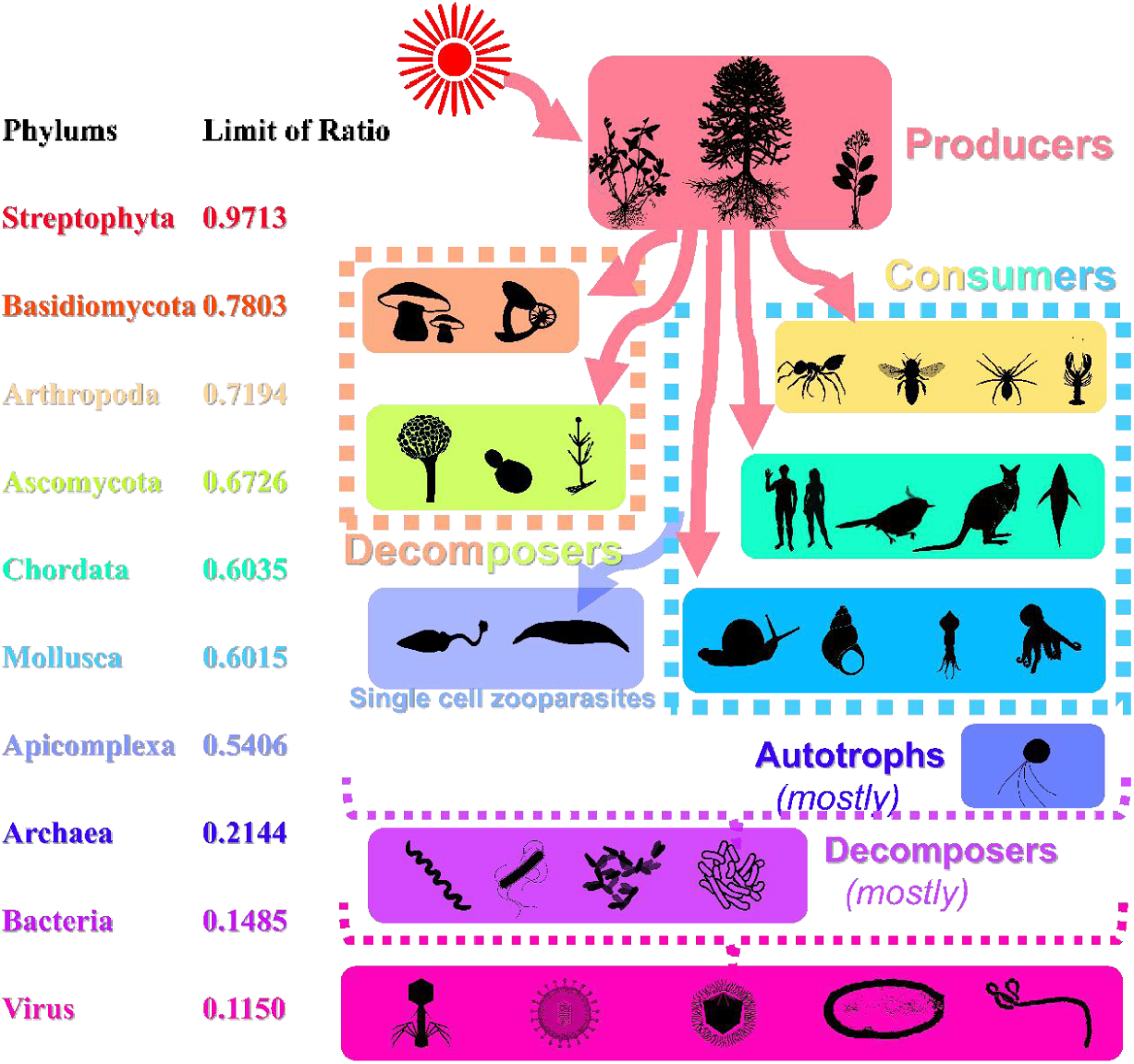
The “Energy Constraint Hypothesis” explaining the proportional limits of non-transcribing genomes across different organisms.

This study has several limitations. First, the significance of the β-value in the linear function remains unclear and may be related to the conserved non-transcribing genome of certain species. Second, the molecular mechanisms behind the linear correlation between non-transcribing region size and genome size are unknown, potentially involving transposon regulation mechanisms. Third, in certain phyla of eukaryotes, the linear relationship exhibits some divergence, suggesting possible further subdivision of phyla, which requires more annotated genomes. Fourth, genome reduction occurs in some parasitic organisms^[23]^, and analyze the gene compositions of parasitic plants/organisms may provide further evidence for the “energy constraint hypothesis,” which also necessitates more annotated genomes. Fifth, I was failed to fit a curve for the nonlinear relationship between coding region size and genome size, which may provide further insights into its biological significance (e.g., the ORF number-genome size curve and Benford distribution)^[24]^.

In conclusion, through quantitative analysis of the three genome components (the non-transcribing genome, non-coding transcribing genome, and coding genome, each with DNA, RNA, and proteins as their respective final functional entities), I observed a linear relationship between genome size and non-transcribing genome size (indicative of proportional genome expansion). I also identified differences in the linear proportional coefficients between different eukaryotic phyla, suggesting that the size and proportion of non-coding regions may be a conserved feature in eukaryotes. I propose the “draft paper” hypothesis for the non-coding genome as raw material for de novo gene formation, and the “energy constraint” hypothesis to explain proportional differences based on the organism’s position in the energy transfer chain of ecosystem.

## Author Approval

The author have seen and approved the manuscript, and that it hasn’t been accepted or published elsewhere.

## Acknowledgment

This study was supported by Professor Zhang Wei and Professor Hu Jinfang from the First Affiliated Hospital of Nanchang University. The downloading of genome annotation files and the writing and running of python scripts for this study were completed by a bioinformatics engineer who wished to remain anonymous.

